# RNApdbee 3.0: A unified web server for comprehensive RNA secondary structure annotation from 3D coordinates

**DOI:** 10.1101/2025.11.07.687208

**Authors:** Jan Pielesiak, Kamil Niznik, Pawel Snioszek, Gabriel Wachowski, Mikolaj Zurawski, Maciej Antczak, Marta Szachniuk, Tomasz Zok

**Affiliations:** Institute of Computing Science, Poznan University of Technology, ul. Jacka Rychlewskiego 1, 61-131, Poznan, Poland; Department of Structural Bioinformatics, Institute of Bioorganic Chemistry PAS, Noskowskiego 12/14, 61-704, Poznan, Poland

**Keywords:** RNA 2D structure annotation, base-pair identification, consensus 2D structure, 2D structure visualization, RNA 3D structure processing

## Abstract

RNApdbee 3.0 (publicly available at https://rnapdbee.cs.put.poznan.pl/) offers an advanced pipeline for comprehensive RNA structural annotation, integrating 2D and 3D data to build detailed nucleotide interaction networks. It classifies base pairs as canonical or noncanonical using the Leontis–Westhof and Saenger schemes and identifies stacking, base-ribose, base-phosphate, and base-triple interactions. The tool handles incomplete or modified residues, marking missing nucleotides and distinguishing noncanonical base pairs for accurate and effective visualization. Results are provided in standard formats – namely, extended dot-bracket notation, BPSEQ, and CT – and in detailed graphical visualizations. RNApdbee decomposes 2D structures into stems, loops, and single strands and offers flexible pseudoknot encoding. Its unified framework addresses inconsistencies across structural data formats by standardizing all inputs to PDBx/mmCIF and integrating seven widely used annotation tools. Finally, RNApdbee ensures reliable, format-independent, and comprehensive RNA structural annotation and interpretation.

## 1 Introduction

Elucidation of RNA 3D structures is essential for understanding diverse functions of this molecule in living organisms. The 2D structure, representing canonical and noncanonical base-pairing patterns, serves as a critical intermediate between the primary sequence and the 3D conformation. Therefore, reliable annotation and visualization of RNA 2D structures derived from experimental or predicted 3D models are key for model validation, comparative analysis, and identification of conserved structural motifs across RNA families.

Over the past two decades, computational tools such as MC-Annotate [12], RNAView [38], 3DNA/DSSR [23], FR3D [31], CompAnnotate [16], Barnaba [7], and BPNnet [29] have been developed to annotate base pairs directly from RNA atomic coordinates. These methods differ in their definitions and detection of canonical and noncanonical interactions, leading to divergent results, particularly for pseudoknotted or otherwise complex RNA architectures [33, 4, 42]. To address these discrepancies, multiple annotation algorithms can be combined within a single framework, allowing systematic comparison and consensus-based interpretation of RNA 2D structures. RNApdbee was introduced as a platform that implements this integrative approach. Version 1.0 [5] provided a web-based platform for extracting 2D structures – including higher-order pseudoknots – from PDB files, encoding them in extended dot-bracket notation (DBN), and visualizing them. RNApdbee 2.0 [43] extended these capabilities with support for additional file formats, expanded pseudoknot characterization, and visualization of canonical, noncanonical, and quadruplex interactions. The system has since incorporated into multiple studies and pipelines for RNA structure analysis and validation [24, 25, 3, 19, 28, 27].

RNApdbee 3.0 presented here is the next-generation platform designed to meet the requirements of modern RNA structural bioinformatics. The system integrates seven base-pair annotation tools and four visualization programs within a format-independent, microservice-based framework. It implements optimized pseudoknot identification and DBN encoding, ensuring reproducible 2D structure annotations. RNApdbee 3.0 also introduces a consensus 2D structure logo that combines multiple annotations, supporting comparative analysis and highlighting conserved interactions. Together, these features provide a comprehensive, scalable, and reproducible resource for interpreting RNA 3D data through consistent 2D annotations, supporting both experimental and computational research. A summary of the evolution of RNApdbee functionality is presented in Table 1. The RNApdbee webserver is publicly available at https://rnapdbee.cs.put.poznan.pl/, and its source code is accessible at https://github.com/rnapdbee/.

**Table 1.**
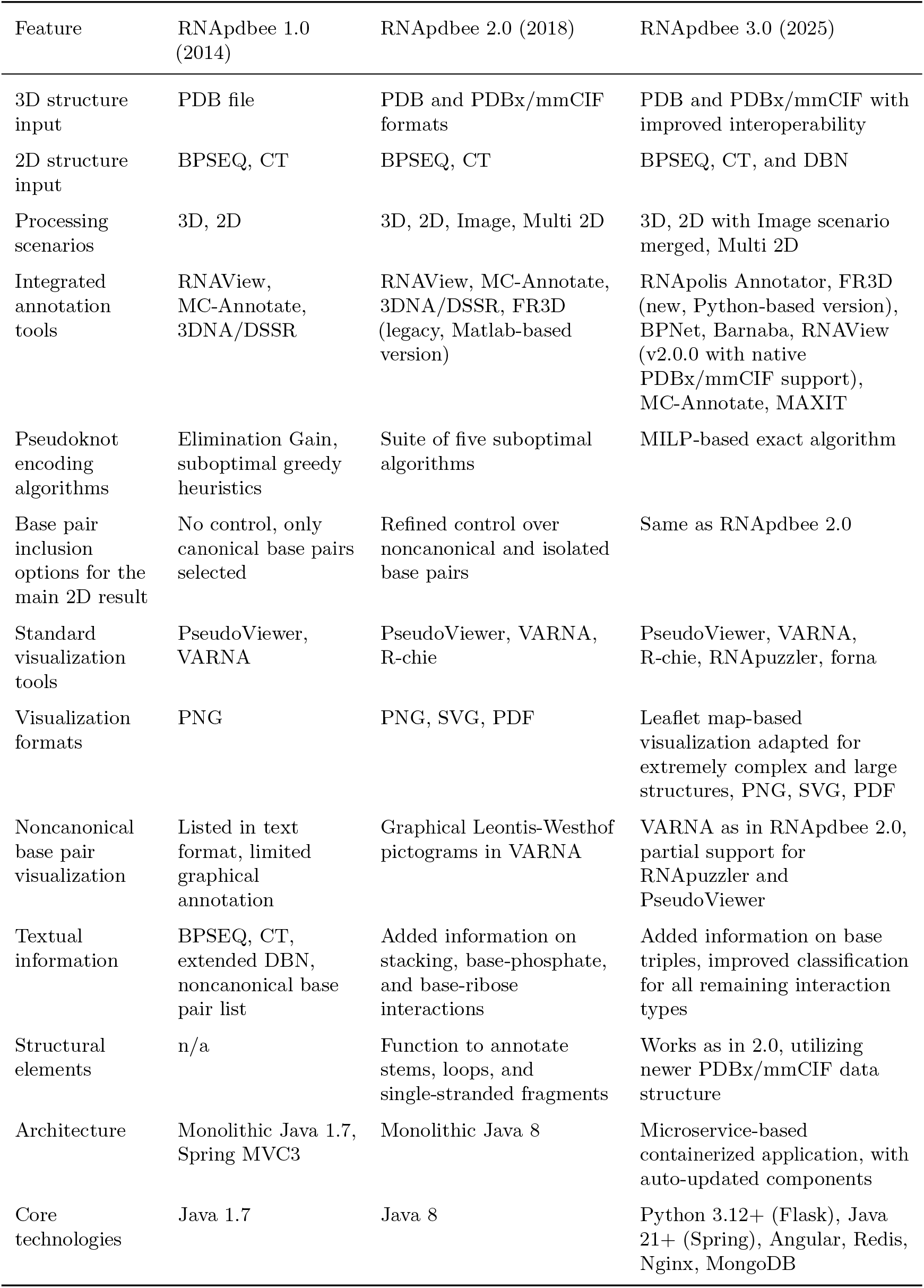
Features of the RNApdbee system across versions 1.0, 2.0, and 3.0.

## 2 Results and Discussion

### 2.1 Advanced RNA structural analysis pipeline

#### 2.1.1 Comprehensive processing of the RNA interaction network

RNApdbee provides advanced analysis of RNA 3D and 2D structures by constructing a comprehensive network of nucleotide interactions. This network includes base pairs and, for 3D input data, also stacking interactions, base–ribose, and base–phosphate contacts. The software classifies all base pairs as canonical (Watson-Crick and G–U wobble) or noncanonical according to the Leontis-Westhof [22] and Saenger [30] schemes. Crucially, RNApdbee applies the Saenger classification universally, even when the input is generated by tools that do not natively provide it. When supported by the underlying data, RNApdbee output also includes stacking topology [17] and detailed base-ribose/base-phosphate classifications [40]. The tool also automatically identifies and classifies base triples based on established nomenclature [2].

#### 2.1.2 Handling of incomplete or modified residues

For the 3D input, unlike conventional tools that process only `ATOM` and `HETATM` coordinate records, RNApdbee accurately represents the complete molecule, even with structural gaps. It incorporates metadata to identify and flag nucleotides with missing atomic coordinates, which are clearly marked with red color in both textual and graphical outputs. The tool also provides robust support for modified residues, ensuring their correct identification and distinct visualization even when standard metadata is missing.

#### 2.1.3 Optimal pseudoknot decomposition

A core innovation in RNApdbee 3.0 is its ability to determine the optimal pseudoknot topology. By replacing previous heuristics, the new version provides a single exact algorithm for encoding pseudoknots in extended dot-bracket notation. This guarantees reproducibility and eliminates ambiguity in structural representation.

In addition, RNApdbee provides two distinct modes for defining structural elements. The default mode uses the complete set of base pairs, including those forming pseudoknots, to provide a direct and faithful representation of the full interaction network. For an H-type pseudoknot formed between a hairpin loop and a single strand, this mode identifies the pseudoknotted pairs as a new stem, thereby redefining the original hairpin and single-stranded regions. Alternatively, a classical mode is available that ignores pseudoknotted pairs during element definition, preserving traditional structural interpretations. The availability of both modes makes RNApdbee a versatile tool, supporting modern, data-driven analysis while remaining compatible with established structural biology workflows.

#### 2.1.4 Versatile output and visualization

Results are delivered in multiple standard formats, including BPSEQ, CT, and an extended dot-bracket notation. The latter explicitly represents pseudoknots of varying orders using paired brackets ([], `{}`, `<>`) and letter cases (`Aa`, `Bb`, etc.) [5, 4]. As mentioned above, it also uses a minus character (-) to denote missing residues. Based on the complete base-pairing network, RNApdbee decomposes the 2D structure into fundamental elements: stems, loops, and single-stranded fragments. This structural decomposition is provided as a detailed textual report that identifies and describes each component strand.

RNApdbee also provides advanced visualization capabilities. We have addressed key limitations of existing tools, which often lack support for pseudoknots or noncanonical pairs. Our custom fork of VARNA (available at https://github.com/tzok/varna-tz) enables direct visualization of pseudoknots in distinct colors, noncanonical interactions using simplified Leontis-Westhof pictograms, and missing residues. For models generated by RNApuzzler and PseudoViewer, RNApdbee post-processes their SVG/EPS output to add a layer rendering color-coded pseudoknots and noncanonical base pairs.

### 2.2 Web Application

RNApdbee 3.0 is available as both a self-deployable resource and a web application. The web application can be accessed in all modern browsers, including mobile browsers. The web interface, written in Angular, is a visual overhaul of the previous version. Based on user feedback, we improved the user experience by adding the *Reanalyze with different parameters* option to the results page. This option allows for a quick change of parameters without requiring the form to be filled over. Moreover, the results of a given request are stored under a unique link that can be copied and saved for later access (the web application provided by the RNApdbee 3.0 team stores the results for 14 days; this can be adjusted if the service is deployed by an individual user). The results, as in the previous version, remain downloadable. Consideration was given to ensuring that the separate elements of the response can be independently downloaded by the user on demand – images can be saved in SVG format after being clicked, and all displayed text can be easily copied to the clipboard. Thanks to the use of the Leaflet package, the high performance and responsiveness of the interface have been maintained, while expanding it with a set of tools that enable convenient and effective visualization preview.

### 2.3 Case studies

#### 2.3.1 Consensus ranking of Zika virus RNA models

To demonstrate how RNApdbee’s integrated framework enables more robust structural comparisons, we performed a computational experiment on 52 candidate models of the Zika virus exonuclease-resistant RNA from the 18th RNA-Puzzles challenge [8]. We downloaded the models and the native target structure from the RNA-Puzzles repository. First, we processed each structure with all seven of RNApdbee’s integrated annotators: Barnaba, BPNet, FR3D, MAXIT, MC-Annotate, RNApolis Annotator, and RNAView. Using all canonical and noncanonical base pairs, we calculated the Interaction Network Fidelity (INF) [26] for each model relative to the target, generating seven distinct rankings.

To synthesize these into a single consensus ranking, we used a bootstrapped statistical procedure. We first computed a composite score for each model using the first principal component (PC1) from a Principal Component Analysis (PCA) [1] of the initial rankings. We then applied bootstrapping with 10,000 iterations to this score to assess statistical confidence, yielding a final median rank and a 95% confidence interval (CI) for each model (Table S1).

This consensus ranking provides a more robust and nuanced assessment than any single analyzer could alone. Our analysis identified a clear group of top-performing models. Models 1-5 from the Das group, models 1-2 from the Chen group, and model 2 from the Dokholyan group consistently occupy the first eight positions. Their narrow 95% CIs confirm their high rank, a finding that remained consistent across the interaction network identification tools used. A similar cluster of models with narrow 95% CIs populates the bottom of the ranking.

In contrast, most models in the middle tier exhibit wide 95% CIs, with their uncertainty often spanning 10-20 ranks. This variability makes it impossible to reliably distinguish their relative performance. The ability to perform this consensus analysis, as enabled by the RNApdbee unified framework, provides researchers with a strong signal to identify and focus on the most promising RNA 3D models for subsequent investigation.

#### 2.3.2 Identification of key interactions in an RNA 3D structure

The self-splicing group I intron from *Tetrahymena thermophila* is a foundational molecule in RNA science [39]. Nearly four decades after its discovery, its complex 3D architecture continues to challenge structural analysis. The crystal structure of its catalytic core, resolved to reveal a base-triple sandwich motif (PDB ID 1X8W), provides a dense network of tertiary interactions that serves as a challenging benchmark for computational annotation tools [14]. Key motifs described in the original publication include the guanosine binding site triples `G414:G264-C311`, `A263:C262-G312`, `A261:A265-U310`, and `A306:C266-G309`. In addition, the authors describe a triple-helical scaffold featuring a ribose-ribose contact (`A104`-`C217`), a base triple (`A105:C216-G257`), and a base-ribose contact (`U106`-`U258`). There is also a domain-connecting fragment comprising a single hydrogen bond pair (`C217`-`C255`), a base triple (`A218:C102-G272`), and base-phosphate contacts between `A256` and `G272`/`U273`. Finally, the authors highlight an interdomain interaction formed by stacked `A324` and `A125`, along with the base triple `A325:G119-U202`.

To assess the consistency of current RNA structure annotation methods, we analyzed the 1X8W structure using RNApdbee, which runs a suite of integrated tools. These programs vary in their capabilities: MAXIT detects only base pairing, Barnaba adds stacking, while others, i.e., MC-Annotate, RNAView, BPNet, FR3D, and RNApolis Annotator, also annotate base-phosphate and base-ribose contacts. To ensure a fair comparison, base triples were identified using RNApdbee’s algorithm rather than the diverse implementations across tools. We note that none of the tested programs are designed to detect ribose-ribose contacts.

Comparative analysis, detailed in Table 2, reveals a stark contrast between the annotation of canonical and noncanonical interactions. Strong consensus exists for canonical Watson-Crick (cWW) base pairs, indicating that their detection is a solved problem. In contrast, the annotation of noncanonical and tertiary contacts is highly inconsistent across the tools. For instance, in the `A263:C262-G312` triple, tools agreed on the Hoogsteen-Sugar (HS) edges but disagreed on the cis/trans orientation. For `A325:G119-U202`, the interaction was variously reported as cSS or cSW. The trans-Hoogsteen/Hoogsteen (tHH) pair in `A306:C266-G309` was identified only by FR3D and RNAView, highlighting a common failure to recognize this geometry. Furthermore, base-phosphate and base-ribose contacts were reported sporadically and were frequently missed entirely.

**Table 2.** RNA interaction detections across seven annotation tools for 12 crucial structural motifs in *Tetrahymena thermophila* ribozyme. Each row represents a specific interaction component (e.g., cWW, tHS, BR) and shows the chains (A, B, C, D) in which each tool identified that component. A dash (-) indicates the tool did not detect that component. Leontis-Westhof nomenclature is used: c = cis, t = trans; W = Watson-Crick edge, H = Hoogsteen edge, S = Sugar edge; BR = base-ribose; BPh = base-phosphate; S = stacking. Some interactions that are sporadically or frequently missed by the listed methods – such as ribose-ribose contacts – may be detected by tools not incorporated into RNApdbee 3.0 (e.g., DSSR).

| Crucial Interaction | Interaction Type | Component | Barnaba | BPNet | FR3D | MAXIT | MC-Annotate | RNApolis Annotator | RNAView |
| --- | --- | --- | --- | --- | --- | --- | --- | --- | --- |
| G414:G264-C311 | Base triple | cWW | A,B,C,D | A,B,C,D | A,B,C,D | A,B,C,D | A,B,C,D | A,B,C,D | A,B,C,D |
|  |  | cHW | A,B,C,D | A,B,C,D | A,B,C,D | - | B,C,D | A,B,C,D | A,B,C,D |
| A263:C262-G312 | Base triple | cWW | B,C,D | B,C,D | B,C,D | B,C,D | B,C,D | B,C,D | B,C,D |
|  |  | cHS | A,B,C,D | - | - | A | - | A,D | - |
|  |  | tHS | - | A,B,C,D | A,B,C,D | - | A,B,C,D | B,C | A,B,C,D |
|  |  | BR | - | C | A,B,C,D | - | A,B,C | A,B,C,D | - |
| A261:A265-U310 | Base triple | cWW | A,B,C,D | A,B,C,D | A,B,C,D | A,B,C,D | A,B,C,D | A,B,C,D | A,B,C,D |
|  |  | tWH | A,B,C,D | C,D | A,B,C,D | - | A,D | B,C,D | A,B,C,D |
|  |  | BPh | - | - | B | - | D | B | - |
| A306:C266-G309 | Base triple | cWW | A,B,C,D | A,B,C,D | A,B,C,D | A,B,C,D | A,B,C,D | A,B,C,D | A,B,C,D |
|  |  | tHH | - | - | A,D | - | - | - | B |
| A104-C217 | Base-ribose | cSS | A,D | B,C,D | A,C,D | - | D | - | - |
|  |  | cWS | - | - | - | - | - | D | - |
|  |  | BR | - | B,D | - | - | A,B,D | A,D | - |
| A105:C216-G257 | Base triple | cWW | A,B,C,D | A,B,C,D | A,B,C,D | A,B,C,D | A,B,C,D | A,B,C,D | A,B,C,D |
|  |  | tSS | B | A,D | A,B,C,D | - | B | B,C,D | A,B,C,D |
|  |  | tWS | - | B | - | - | - | - | - |
|  |  | BR | - | B | C,D | - | B | B,C,D | - |
| U106-U258 | Ribose-ribose | - | - | - | - | - | - | - | - |
| C217-C255 | Base pair (1H) | tWH | B,C,D | D | C,D | - | - | C,D | B,C,D |
|  |  | tWW | - | - | - | B | - | - | - |
| A218:C102-G272 | Base triple | cWW | A,B,D | A,B,D | A,B,D | A,B,D | A,B,D | A,B,D | A,B,D |
|  |  | cSS | A,B | A,B | A,B | - | B | A,B | - |
| A256-G272 | Base-phosphate | BPh | - | A,B,C,D | - | - | - | C,D | - |
|  |  | BR | - | - | A,B,D | - | - | B | - |
| A125-A324 | Stacking | S | B,D | D | B,D | - | B,D | B,D | - |
|  |  | BR | - | A | B | - | A | A,B | - |
| A325:G119-U202 | Base triple | cWW | A,B,C | A,B,D | A,B,C,D | A,B,C,D | A,B,C | A,B,C,D | A,B,C,D |
|  |  | cSS | B | A | A,B | - | A | - | B |
|  |  | cSW | A | B | - | - | - | A,B | - |
|  |  | S | - | - | D | - | - | - | - |
|  |  | BR | - | A,B | - | - | A,B,D | A,B | A |

This divergence in performance reveals distinct annotation profiles among the tools. MAXIT is consistently the most conservative, reporting only high-confidence interactions. At the other extreme, FR3D and RNApolis Annotator provide the most exhaustive annotations, possibly at the risk of including ambiguous contacts. Barnaba, BPNet, and RNAView occupy a middle ground – they detect more noncanonical interactions than MAXIT but are less comprehensive than MC-Annotate, FR3D, and RNApolis Annotator, particularly for base-ribose and base-phosphate contacts.

These findings underscore that no single tool currently serves as a universal gold standard for annotating noncanonical RNA interactions. The choice of tool should therefore depend on the research objective: MAXIT is suitable for a high-confidence overview, whereas FR3D or RNApolis Annotator are better for in-depth, exploratory analysis. Crucially, the results demonstrate that human oversight remains essential, as no single program can be trusted to produce a perfectly accurate annotation. A consensus-based approach that integrates outputs from multiple tools offers a pragmatic path forward. Ultimately, the RNA structural bioinformatics field would benefit immensely from the establishment of standardized geometric criteria and a community-accepted reference implementation to improve the reproducibility and reliability of its analyses.

## 3 Materials and Methods

### 3.1 System architecture

The RNApdbee backend is built on a resilient microservice architecture designed for high availability and performance. To manage demanding workloads and ensure stability, the system incorporates component replication, dynamic load-balancing, and automated recovery mechanisms based on service health checks and restart policies.

The architecture employs a dual-storage strategy for data management. Redis, a high-performance, in-memory cache, provides low-latency access to intermediate analysis data, such as raw results from external tools that are likely to be reused. Final run outputs, including all annotations and visualizations, are persistently stored in MongoDB, a NoSQL database optimized for complex, JSON-like documents.

### 3.2 Standardized processing pipeline

The analysis of RNA structural data is hindered by a fragmented landscape of data formats and incompatible tools. RNApdbee addresses this by providing a robust, unified platform (Figure 1). The first step is adopting PDBx/mmCIF as the canonical internal data representation. Structures provided in the legacy PDB format are converted losslessly. This standardized data then undergoes several preprocessing steps, including correcting missing occupancy data, which can cause certain tools to fail. To accommodate tools that do not support the PDBx/mmCIF format, RNApdbee temporarily converts the processed data back into the required PDB format. This back-conversion intelligently remaps chain identifiers and renumbers residues, using the insertion code column as needed. After the tool generates its output, the results are converted back, ensuring that all nucleotide identifiers in the final output precisely match those in the original input file. This allows any of the seven integrated tools to be used with any input format.

**Figure 1.**
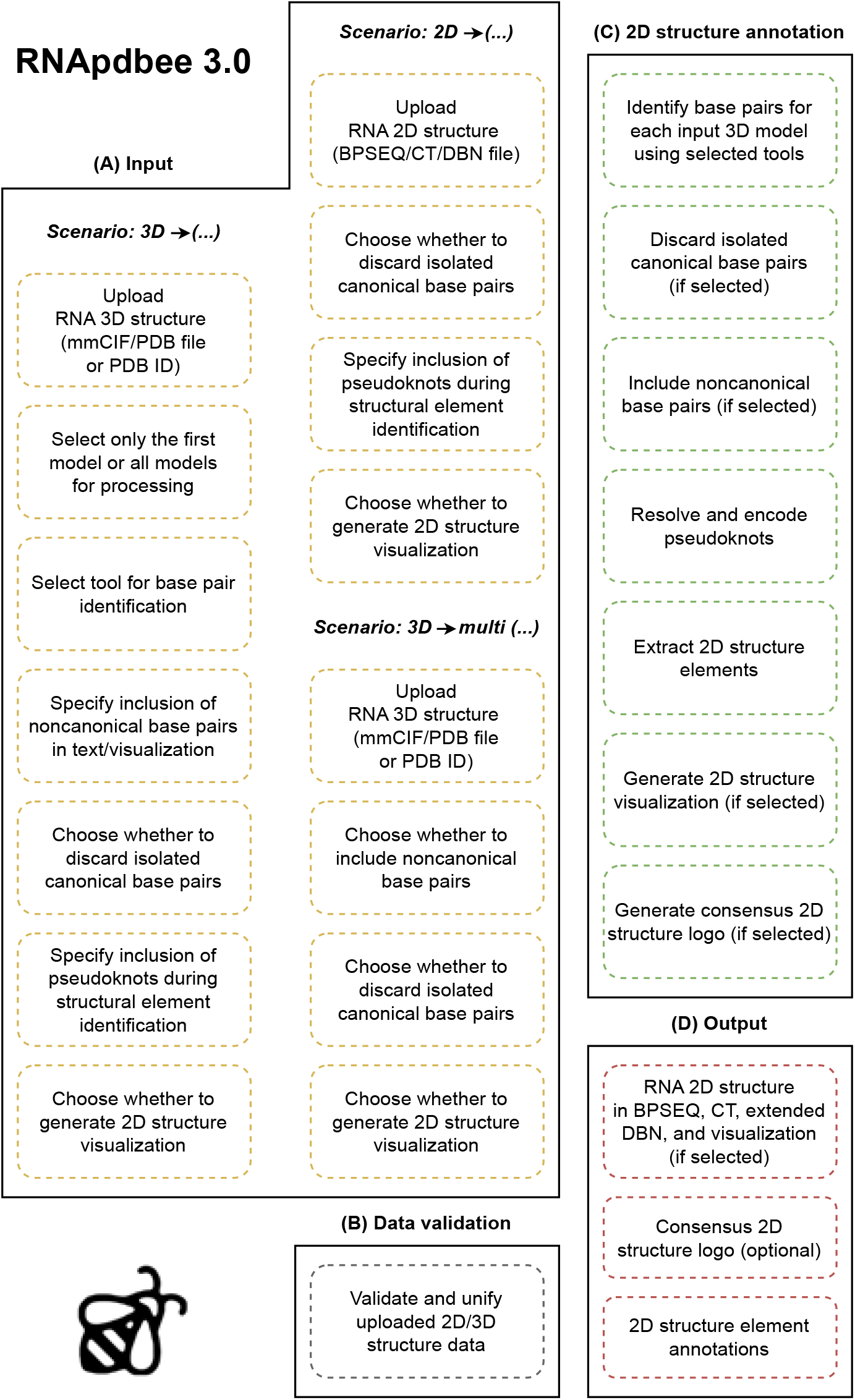
Workflow of RNApdbee 3.0.

RNApdbee integrates critical metadata to provide a more complete representation of the RNA. The software parses `REMARK` lines from PDB files and information on unobserved residues from PDBx/mmCIF files to identify nucleotides with missing atomic coordinates. This information is incorporated directly into the output. For instance, a missing residue is represented by a minus sign (-) in an extended dot-bracket notation and is visually highlighted in the generated diagrams. RNApdbee also provides robust support for modified residues through a double-attempt approach. It first attempts to identify modifications by parsing header information in both PDB and PDBx/mmCIF formats. When this metadata is incomplete or absent, RNApdbee employs a custom modification detector. This algorithm identifies the most closely related canonical nucleotide to ensure accurate sequence representation. To distinguish these from standard nucleotides, modified residues are consistently displayed in lower-case letters in both textual and graphical outputs.

### 3.3 Resolving ambiguities

The unification framework also resolves critical ambiguities in residue identification. The PDBx/mmCIF format uses two parallel schemes: `auth` columns represent author-provided numbering (which can be non-sequential, negative, or use insertion codes), while `label` columns enforce a strictly sequential, 1-based index. Because different tools may rely on either scheme, RNApdbee tracks both `auth` and `label` identifiers for every nucleotide. This guarantees that each residue is unambiguously identified, regardless of the conventions used by a specific analysis tool.

Finally, RNApdbee standardizes and enriches its output. It includes a classification module that uses sequence information and the Leontis-Westhof classification of a base pair to infer the corresponding Saenger’s notation, if applicable. As a result, users receive Saenger classifications even if the underlying analysis tool does not natively provide them, ensuring a consistent and comprehensive analysis.

### 3.4 Conversion between PDBx/mmCIF and PDB formats

RCSB MAXIT [11] was developed to format and validate macromolecular structure data in PDB and PDBx/mmCIF formats, ensuring conformance with data standards and dictionary definitions during deposition of 3D structures to the Protein Data Bank. We used this tool to convert 3D coordinates between PDBx/mmCIF and PDB formats and vice versa. All RNA 3D structures identified by the user’s PDB IDs are retrieved and cached from the Protein Data Bank [6]. The Protein Data Bank is the first open-access resource for experimentally determined macromolecular structures, established in 1971. RNApdbee 3.0 uses the BioCommons library [41] to transform the interaction network derived from 3D structure into a 2D structure in BPSEQ format supplemented with noncanonical pairings.

### 3.5 Base-pair identification

A cornerstone of the RNApdbee 3.0 workflow is its expanded suite of tools for identifying and classifying nucleobase interactions. To guarantee long-term stability and unrestricted distribution of our Docker-based system, we replaced 3DNA/DSSR [23] with alternatives that offer permissive licenses. Specifically, current licensing restrictions for 3DNA and DSSR prevent their redistribution in containerized environments and mandate user-specific validation that is incompatible with our open, login-free web server architecture. The new workflow retains RNAView [38], MC-Annotate [12], and FR3D [31], while integrating BPNet [29], Barnaba [7], and RNApolis Annotator [35, 42]. In addition, the user can now opt to use data from `ndb_struct_na_base_pair` columns from the PDBx/mmCIF produced by MAXIT [11] as the source of base pairing information. This diverse collection enables users to perform extensive comparative analyses of annotation results.

We selected RNApolis Annotator as the new default for the RNApdbee 3.0 processing pipeline due to its modern architecture and continuous development. Implemented in Python and available on PyPI, it supports both PDB and PDBx/mmCIF file formats and can process multiple models concurrently. The tool identifies base-base, base-phosphate group, base-ribose, and stacking interactions.

The other integrated tools offer distinct capabilities. FR3D, notable for its Python implementation and active development, is a widely used tool for base-pairing exploration, as evidenced by its adoption across platforms such as the Nucleic Acid Knowledgebase [21]. It only processes PDBx/mmCIF format and identifies base-base interactions (Leontis-Westhof classification only) and stacking interactions. BPNet handles both PDB and PDBx/mmCIF files, allows chain selection, and can exclude heteroatoms. It annotates base-base (Leontis-Westhof only), base-phosphate, base-ribose, and stacking interactions, and provides primary and 2D structures in FASTA, DBN, and BPSEQ formats. Barnaba is designed to analyze RNA structures and molecular dynamics trajectories, with motif searching and SVG visualization capabilities. However, its annotations are limited to base-base (Leontis-Westhof) and stacking interactions, and it does not support the PDBx/mmCIF format.

RNApdbee also includes and extends the legacy tools. MC-Annotate supports both Leontis-Westhof and Saenger classifications for base-base interactions and also identifies base-phosphate, base-ribose, and stacking interactions. It allows analysis of specific residue ranges but is restricted to the legacy PDB format. RNAView identifies a similar range of interactions – base-base (Leontis-Westhof and Saenger), base-ribose, and base-phosphate – and can generate PostScript visualizations. It can process a specified chain or multiple PDB files in a single run, and its newest version now supports the PDBx/mmCIF format.

The base-triple detection pipeline in RNApdbee 3.0 runs in three steps. First, a user-selected annotation tool generates a list of base pairs. Next, the system identifies base triple candidates from this list. Finally, each candidate undergoes geometric validation. The pipeline depends entirely on the chosen base pair analyzer. As a result, final base triple identification varies depending on the tool used, such as FR3D, Barnaba, or RNAView. If the analyzer misses constituent base pairs, those nucleotides cannot be forwarded as candidates.

To ensure high-fidelity detection of these higher-order interactions, we implemented a rigorous geometric validation step for the collected base triples. It is not enough to merely detect a nucleotide simultaneously forming two distinct base pairs, as this approach can yield false positives if the bases are not spatially aligned. RNApdbee 3.0 addresses this by enforcing coplanarity. For every candidate triple, we compute a global best-fit plane derived from the core ring atoms of all three residues. A triple is accepted only if it satisfies two geometric criteria: (1) the centroid of each nucleobase must lie within a maximum distance of 0.2 Å from the global plane, and (2) the normal vector of each base must align with the global plane normal within a maximum deviation of 25°. Candidates failing these checks are filtered out of the main result table and reported separately to maintain structural accuracy.

### 3.6 2D structure visualization

Effective visualization of RNA 2D structure requires rendering complex features such as pseudoknots, multistranded complexes, and noncanonical base pairs. However, existing standalone tools often lack comprehensive support for these elements. To address this gap, RNApdbee 3.0 integrates and extends a suite of established visualization programs to provide a complete visualization solution: VARNA [10], PseudoViewer [9], R-chie [20], and RNApuzzler [37]. In addition, RNApdbee generates dynamic links to forna [18] – an interactive, in-the-browser visualization tool.

Among these, the RNApuzzler tool [37] stands out thanks to its performance and the aesthetic quality of its output. As one of the algorithms within the ViennaRNA package’s RNAplot program, RNApuzzler processes sequence and 2D structure data to generate overlap-free diagrams in Encapsulated PostScript format. We leveraged this flexible output to significantly expand its capabilities. Our implementation introduces clear rendering for noncanonical base pairs, pseudoknots, residues with missing coordinates, and multi-stranded structures. We also modified the tool’s default color palette for improved clarity.

Our integration of PseudoViewer [9], a tool available as a web application, web service, and standalone program, also provides new functionality. While PseudoViewer natively supports interactive SVG diagrams of pseudoknotted and multistranded structures, RNApdbee 3.0 introduces several key improvements. We modified the pseudoknot drawing style, implemented mechanisms to display noncanonical base pairs and missing residues, and optimized the SVG file to minimize its size.

For arc diagram representations, RNApdbee 3.0 incorporates the R-chie package [20]. Natively, R-chie renders pseudoknots, covariances, and comparative visualizations of two structures into PDF format. Our integration enhances this by implementing a consistent color palette for displaying pseudoknots of successive orders and by providing conversion from native PDF output to SVG.

The visualization suite is completed by VARNA [10], a highly versatile tool that is selected as the default in RNApdbee 3.0. VARNA’s strengths include support for a wide range of input formats (BPSEQ, CT, and extended DBN) and multiple output formats (EPS, SVG, JPG, and PNG). Notably, VARNA is the only integrated tool that offers out-of-the-box support for marking noncanonical base pairs according to the Leontis-Westhof classification, making it a valuable complementary component of the RNApdbee 3.0 toolkit. To meet our specific requirements, we implemented substantial customizations and new features directly into a fork of the original project, available at https://github.com/tzok/varna-tz.

### 3.7 Exact pseudoknot encoding

A key enhancement in RNApdbee 3.0 is the refinement of pseudoknot encoding in the extended dot-bracket notation. This version replaces the heuristics-based approach of version 2.0 with an optimal algorithm that solves the Pseudoknot Order Assignment Problem (POA) [42]. While the previous version generated several suboptimal solutions that required user evaluation, version 3.0 provides a single, exact algorithm. The new algorithm achieves this by reducing the POA to a Mixed-Integer Linear Programming (MILP) problem, which is then solved efficiently using the HiGHS solver [15]. The solver operates on two strict criteria: it primarily minimizes the overall pseudoknot order, and secondarily ensures that the maximum number of pairs are retained in the non-pseudoknotted (nested) structure before populating higher orders. It allows users to obtain unambiguous, optimal, and repeatable solutions for two criteria. On the other hand, it has simplified the application interface, as there is no longer a need to select algorithms for classifying pseudoknots.

### 3.8 Consensus 2D structure construction

The seven base-pair identification tools integrated into RNApdbee 3.0 can produce significantly different results for a single RNA 3D structure, with discrepancies particularly pronounced for noncanonical base pairs, long-range interactions, and pseudoknots. To synthesize these diverse annotations, we developed a procedure that generates a consensus 2D structure, visualized in a format inspired by sequence logos [32]. This representation concisely displays the diversity and prevalence of structural annotations at each nucleotide position on a single diagram. The logo is constructed by aligning 2D structures using the Python-based Logomaker package [36]. In our visualization, unpaired nucleotides and missing residues are denoted by “U” and “Z”, respectively, while pseudoknots are colored using the established RNApdbee palette for visual consistency. It offers a significant advantage over individual tools when analyzing complex 2D structures such as SARS-CoV-2 UTRs [13], which also contain pseudoknots.

However, we emphasize that this consensus visualization should be interpreted with a semi-automatic approach rather than as absolute ground truth. Due to the heavy imbalance of interaction types in nature, where canonical cWW pairs dominate [34], and the potential for correlated errors among annotation tools, high agreement levels can occasionally mask incorrect interactions. This is most critical for rare non-canonical base pairs. Therefore, we recommend treating these consensus annotations as structural hypotheses that warrant verification against experimental quality indicators, such as B-factors or map resolution.

## 4 Conclusions

RNApdbee addresses the critical challenge of fragmentation and inconsistency in RNA structural bioinformatics. Its unification framework resolves the widespread misalignment between specialized tools and disparate data formats, enabling legacy software to process modern files and extending the capabilities of existing programs. Key technical innovations include an optimal algorithm for encoding pseudoknotted structures in extended dot-bracket notation and the novel use of Sequence Logos to represent structural consensus. These features are delivered through a user-friendly and efficient interface, ensuring broad accessibility.

To demonstrate its practical utility, we applied RNApdbee to two distinct challenges. We successfully used the tool to rank computational models submitted to the RNA-Puzzles 18th challenge using a consensus-based approach. Furthermore, we analyzed the crucial yet elusive tertiary interactions within the *Tetrahymena thermophila* ribozyme, showing how RNApdbee simplifies the identification and characterization of complex structural features. These applications underscore the tool’s immediate impact on real-world research problems. Taken together, the advanced capabilities and demonstrated performance of RNApdbee 3.0 position it to become a standard tool for RNA structural analysis.

## 5 CRediT authorship contribution statement

**Jan Pielesiak:** Software, Writing - original draft. **Kamil Niznik**: Software, Visualization. **Pawel Snioszek**: Software, Validation. **Gabriel Wachowski**: Software, Visualization, **Mikolaj Zurawski**: Software, Validation. **Maciej Antczak**: Conceptualization, Methodology, Software, Writing - original draft. **Marta Szachniuk**: Conceptualization, Methodology, Writing - original draft. **Tomasz Zok**: Conceptualization, Funding acquisition, Methodology, Project administration, Software, Writing - original draft.

## Acknowledgements

This work is supported by funds from the National Science Center, Poland [2023/51/D/ST6/01207 to TZ] and statutory funds of the Institute of Computing Science, Poznan University of Technology. The authors are grateful to Dr Mariusz Popenda and Dr Joanna Sarzynska for their valuable suggestions that contributed to the improvement of RNApdbee and the development of version 3.0.

## References

[1] H. Abdi and L. J. Williams. “Principal component analysis”. In: Wiley interdisciplinary reviews: computational statistics 2.4 (2010), pp. 433–459. DOI: 10.1002/wics.101.

[2] S. Abu Almakarem et al. “Comprehensive Survey and Geometric Classification of Base Triples in RNA Structures”. In: Nucleic Acids Res 40.4 (2012), pp. 1407–1423. DOI: 10.1093/nar/gkr810.

[3] B. Adamczyk et al. “WebTetrado: a webserver to explore quadruplexes in nucleic acid 3D structures”. In: Nucleic Acids Res 51.W1 (2023), W607–W612. DOI: 10.1093/nar/gkad346.

[4] M. Antczak et al. “New algorithms to represent complex pseudoknotted RNA structures in dot-bracket notation”. In: Bioinformatics 34.8 (2017), pp. 1304–1312. DOI: 10.1093/bioinformatics/btx783.

[5] M. Antczak et al. “RNApdbee — a webserver to derive secondary structures from pdb files of knotted and unknotted RNAs”. In: Nucleic Acids Res 42.W1 (2014), W368–W372. DOI: 10.1093/nar/gku330.

[6] H. M. Berman et al. “The Protein Data Bank”. In: Nucleic Acids Res 28.1 (2000), pp. 235–242. DOI: 10.1093/nar/28.1.235.

[7] S. Bottaro et al. “Barnaba: software for analysis of nucleic acid structures and trajectories”. In: RNA 25.2 (2019), pp. 219–231. DOI: 10.1261/rna.067678.118.

[8] F. Bu et al. “RNA-Puzzles Round V: Blind Predictions of 23 RNA Structures”. In: Nat Methods 22.2 (2025), pp. 399–411. DOI: 10.1038/s41592-024-02543-9.

[9] Y. Byun and K. Han. “PseudoViewer: web application and web service for visualizing RNA pseudoknots and secondary structures”. In: Nucleic Acids Res 34.suppl_2 (2006), W416–W422. DOI: 10.1093/nar/gkl210.

[10] K. Darty, A. Denise, and Y. Ponty. “VARNA: Interactive drawing and editing of the RNA secondary structure”. In: Bioinformatics 25.15 (2009), p. 1974. DOI: 10.1093/bioinformatics/btp250.

[11] Z. Feng et al. “MAXIT: macromolecular exchange and input tool”. In: Rutgers University, New Brunswick (1998).

[12] P. Gendron, S. Lemieux, and F. Major. “Quantitative analysis of nucleic acid three-dimensional structures”. In: J Mol Biol 308.5 (2001), pp. 919–936. DOI: 10.1006/jmbi.2001.4626.

[13] J. Gumna et al. “Computational pipeline for reference-free comparative analysis of RNA 3D structures applied to SARS-CoV-2 UTR models”. In: Int J Mol Sci 23.17 (2022), p. 9630. DOI: 10.3390/ijms23179630.

[14] F. Guo, A. R. Gooding, and T. R. Cech. “Structure of the Tetrahymena Ribozyme”. In: Mol Cell 16.3 (2004), pp. 351–362. DOI: 10.1016/j.molcel.2004.10.003.

[15] Q. Huangfu and J. A. J. Hall. “Parallelizing the Dual Revised Simplex Method”. In: Math Program Comput 10.1 (2018), pp. 119–142. DOI: 10.1007/s12532-017-0130-5.

[16] S. Islam, P. Ge, and S. Zhang. “CompAnnotate: a comparative approach to annotate base-pairing interactions in RNA 3D structures”. In: Nucleic Acids Res 45.14 (2017), e136–e136. DOI: 10.1093/nar/gkx538.

[17] A. Jhunjhunwala et al. “On the Nature of Nucleobase Stacking in RNA: A Comprehensive Survey of Its Structural Variability and a Systematic Classification of Associated Interactions”. In: J Chem Inf Model 61.3 (2021), pp. 1470–1480. DOI: 10.1021/acs.jcim.0c01225.

[18] P. Kerpedjiev, S. Hammer, and I. L. Hofacker. “Forna (Force-Directed RNA): Simple and Effective Online RNA Secondary Structure Diagrams”. In: Bioinformatics 31.20 (2015), pp. 3377–3379. DOI: 10.1093/bioinformatics/btv372.

[19] M. Kulkarni et al. “Comparative analysis of RNA secondary structure accuracy on predicted RNA 3D models”. In: PloS ONE 18.9 (2023), e0290907. DOI: 10.1371/journal.pone.0290907.

[20] D. Lai et al. “R-CHIE: a web server and R package for visualizing RNA secondary structures”. In: Nucleic Acids Res 40.12 (2012), e95–e95. DOI: 10.1093/nar/gks241.

[21] C. L. Lawson et al. “The Nucleic Acid Knowledgebase: a new portal for 3D structural information about nucleic acids”. In: Nucleic Acids Res 52.D1 (2024), pp. D245–D254. DOI: 10.1093/nar/gkad957.

[22] N. B. Leontis and E. Westhof. “Geometric nomenclature and classification of RNA base pairs”. In: RNA 7.4 (2001), pp. 499–512. DOI: 10.1017/S1355838201002515.

[23] X.-J. Lu, H. J. Bussemaker, and W. K. Olson. “DSSR: an integrated software tool for dissecting the spatial structure of RNA”. In: Nucleic Acids Res 43.21 (2015), e142–e142. DOI: 10.1093/nar/gkv716.

[24] K. Luwanski et al. “RNAspider: a webserver to analyze entanglements in RNA 3D structures”. In: Nucleic Acids Res 50.W1 (2022), W663–W669. DOI: 10.1093/nar/gkac218.

[25] S. S. Nasaev et al. “AliNA – a deep learning program for RNA secondary structure prediction”. In: Mol Inform 42.12 (2023), e202300113. DOI: 10.1002/minf.202300113.

[26] M. Parisien et al. “New Metrics for Comparing and Assessing Discrepancies between RNA 3D Structures and Models”. In: RNA 15.10 (2009), pp. 1875–1885. DOI: 10.1261/rna.1700409.

[27] J. Pielesiak et al. “RNAtive to recognize native-like structure in a set of RNA 3D models”. In: Bioinformatics 41.11 (2025), btaf601. DOI: 10.1093/bioinformatics/btaf601.

[28] M. Quadrini et al. “TARNAS: a software tool for abstracting and translating RNA secondary structures”. In: Int J Mol Sci 26.12 (2025), p. 5728. DOI: 10.3390/ijms26125728.

[29] P. Roy and D. Bhattacharyya. “Contact networks in RNA: a structural bioinformatics study with a new tool”. In: J Comput Aided Mol Des 36.2 (2022), pp. 131–140. DOI: 10.1007/s10822-021-00438-x.

[30] W. Saenger. Principles of Nucleic Acid Structure. Springer Advanced Texts in Chemistry. New York, NY: Springer New York, 1984. ISBN: 978-0-387-90761-1 978-1-4612-5190-3. DOI: 10.1007/978-1-4612-5190-3.

[31] M. Sarver et al. “FR3D: finding local and composite recurrent structural motifs in RNA 3D structures”. In: J Math Biol 56.1 (2008), pp. 215–252. DOI: 10.1007/s00285-007-0110-x.

[32] T. D. Schneider and R. M. Stephens. “Sequence logos: a new way to display consensus sequences”. In: Nucleic Acids Res 18.20 (1990), pp. 6097–6100. DOI: 10.1093/nar/18.20.6097.

[33] S. Smit et al. “From knotted to nested RNA structures: a variety of computational methods for pseudoknot removal”. In: RNA 14.3 (2008), pp. 410–416. DOI: 10.1261/rna.881308.

[34] J. Stombaugh et al. “Frequency and Isostericity of RNA Base Pairs”. In: Nucleic Acids Res 37.7 (2009), pp. 2294–2312. DOI: 10.1093/nar/gkp011.

[35] M. Szachniuk. “RNApolis: computational platform for RNA structure analysis”. In: Found Comput Decis Sci 44 (2019), pp. 241–257. DOI: 10.2478/fcds-2019-0012.

[36] A. Tareen and J. B. Kinney. “Logomaker: Beautiful Sequence Logos in Python”. In:Bioinformatics 36.7 (2020), pp. 2272–2274. DOI: 10.1093/bioinformatics/btz921.

[37] D. Wiegreffe et al. “RNApuzzler: efficient outerplanar drawing of RNA-secondary structures”. In: Bioinformatics 35.8 (2019), pp. 1342–1349. DOI: 10.1093/bioinformatics/bty817.

[38] H. Yang et al. “Tools for the automatic identification and classification of RNA base pairs”. In: Nucleic Acids Res 31.13 (2003), pp. 3450–3460. DOI: 10.1093/nar/gkg529.

[39] J. Zaug, M. D. Been, and T. R. Cech. “The Tetrahymena Ribozyme Acts like an RNA Restriction Endonuclease”. In: Nat 324.6096 (1986), pp. 429–433. DOI: 10.1038/324429a0.

[40] L. Zirbel et al. “Classification and Energetics of the Base-Phosphate Interactions in RNA”. In: Nucleic Acids Res 37.15 (2009), pp. 4898–4918. DOI: 10.1093/nar/gkp468.

[41] T. Zok. “BioCommons: a robust Java library for RNA structural bioinformatics”. In: Bioinformatics 37.17 (2021), pp. 2766–2767. DOI: 10.1093/bioinformatics/btab069.

[42] T. Zok et al. “New models and algorithms for RNA pseudoknot order assignment”. In: Int J Appl Math Comput Sci 30.2 (2020), pp. 315–324. DOI: 10.34768/amcs-2020-0024.

[43] T. Zok et al. “RNApdbee 2.0: multifunctional tool for RNA structure annotation”. In: Nucleic Acids Res 46.W1 (2018), W30–W35. DOI: 10.1093/nar/gky314.

