## Supplementary Materials for "RNApdbee 3.0: A unified web server for comprehensive RNA secondary structure annotation from 3D coordinates" (from another version of this preprint: version 2, 2026-02-14)

Jan Pielesiak[1] Kamil Niznik[1] Pawel Snioszek[1]
Gabriel Wachowski[1] Mikolaj Zurawski[1] Maciej Antczak[1,2]
Marta Szachniuk[1,2] Tomasz Zok[1]

Table S1: Rankings of models submitted to the 18th RNA-Puzzles challenge, presented as medians and 95% confidence intervals (CIs). The rankings are based on a composite INF score computed using all base-pair analyzers integrated within the RNApdbee platform.

| Group | Model | Median rank | Rank 95% CI lower | Rank 95% CI upper |
|---|---|---|---|---|
| Das | 1 | 1 | 1 | 2 |
| Das | 3 | 2 | 1 | 5 |
| Das | 2 | 3 | 2 | 4 |
| Chen | 1 | 4 | 2 | 5 |
| Chen | 2 | 5 | 3 | 6 |
| Das | 4 | 6 | 5 | 6 |
| Das | 5 | 7 | 7 | 7 |
| Dokholyan | 2 | 8 | 8 | 9 |
| RNAComposer | 2 | 9 | 8 | 13 |
| Dokholyan | 1 | 10 | 9 | 12 |
| Dokholyan | 3 | 12 | 9 | 21 |
| 3dRNA | 3 | 13 | 10 | 20 |
| Chen | 3 | 14 | 10 | 29 |
| RW3D | 4 | 15 | 10 | 28 |

*

[1] Institute of Computing Science, Poznan University of Technology, Piotrowo 2, 60-965, Poznan, Poland
[2] Institute of Bioorganic Chemistry, Polish Academy of Sciences, Noskowskiego 12/14, 61-704, Poznan, Poland

| Group | Model | Median rank | Rank 95% CI lower | Rank 95% CI upper |
|---|---|---|---|---|
| RW3D | 9 | 15 | 10 | 23 |
| RW3D | 7 | 15 | 10 | 23 |
| Ding | 2 | 16 | 12 | 22 |
| YagoubAli | 1 | 18 | 14 | 26 |
| RW3D | 6 | 19 | 12 | 26 |
| RNAComposer | 1 | 19 | 13 | 27 |
| RW3D | 10 | 21 | 15 | 30 |
| RNAComposer | 4 | 23 | 14 | 32 |
| 3dRNA | 4 | 24 | 19 | 31 |
| RW3D | 5 | 26 | 20 | 31 |
| 3dRNA | 1 | 26 | 21 | 30 |
| RW3D | 8 | 26 | 17 | 33 |
| RW3D | 2 | 27 | 17 | 34 |
| 3dRNA | 2 | 27 | 17 | 33 |
| RW3D | 1 | 28 | 18 | 34 |
| Chen | 4 | 30 | 13 | 35 |
| RW3D | 3 | 30 | 24 | 34 |
| Ding | 5 | 31 | 24 | 34 |
| 3dRNA | 5 | 32 | 26 | 35 |
| Chen | 5 | 33 | 25 | 35 |
| SimRNA | 2 | 35 | 35 | 36 |
| RNAComposer | 5 | 36 | 22 | 48 |
| Ding | 3 | 37 | 36 | 39 |
| RNAComposer | 3 | 38 | 36 | 42 |
| Ding | 1 | 39 | 37 | 42 |
| Lee | 2 | 39 | 37 | 41 |
| LeeASmodel | 2 | 41 | 37 | 42 |
| Lee | 5 | 42 | 39 | 44 |
| LeeASmodel | 5 | 43 | 41 | 44 |
| Lee | 4 | 44 | 42 | 45 |
| LeeASmodel | 4 | 45 | 44 | 46 |
| Lee | 1 | 46 | 43 | 47 |
| LeeASmodel | 1 | 47 | 45 | 47 |
| SimRNA | 3 | 48 | 47 | 48 |
| SimRNA | 1 | 49 | 49 | 50 |
| Lee | 3 | 50 | 49 | 51 |
| LeeASmodel | 3 | 51 | 50 | 51 |
| Ding | 4 | 52 | 52 | 52 |
